# Altered action potential waveform and shorter axonal initial segment in hiPSC-derived motor neurons with mutations in *VRK1*

**DOI:** 10.1101/2021.04.01.438138

**Authors:** Rémi Bos, Khalil Rihan, Lara El-Bazzal, Nathalie Bernard-Marissal, Patrice Quintana, Rosette Jabbour, André Mégarbané, Marc Bartoli, Frédéric Brocard, Valérie Delague

**Author notes:** **Correspondence to:** Valérie Delague, U 1251, Marseille Medical Genetics, Faculté de Médecine de la Timone, 27 bd Jean Moulin, 13385 Marseille cedex 05. These authors contributed equally (Rémi Bos, Khalil Rihan). These authors contributed equally (Frédéric Brocard, Valérie Delague).

## Abstract

We recently described new pathogenic variants in *VRK1*, in patients affected with distal Hereditary Motor Neuropathy associated with upper motor neurons signs. Specifically, we provided evidences that hiPSC-derived Motor Neurons (hiPSC-MN) from these patients display Cajal bodies (CBs) disassembly and defects in neurite outgrowth and branching.

We here focused on the Axonal Initial Segment (AIS) and the related firing properties of hiPSC-MNs from these patients. We found that the patient’s Action Potential (AP) was smaller in amplitude, larger in duration, and displayed a more depolarized threshold while the firing patterns were not altered. These alterations were accompanied by a decrease in the AIS length measured in patients’ hiPSC-MNs. These data indicate that mutations in VRK1 impact the AP waveform and the AIS organization in MNs and may ultimately lead to the related motor neuron disease.

**Highlights:**

- hiPSC-MNs are functional and sustain firing patterns, typical of spinal MNs
- hiPSC-MNs from patients with VRK1 mutations have altered Action Potential
- Axonal Initial Segment is shorter in hiPSC-MNs from patients with mutated VRK1
- hiPSC-MNs are a useful platform to study Motor Inherited Peripheral Neuropathies

**eTOC Blurb:** In human spinal Motor Neurons derived from induced Pluripotent Stem Cells from patients with VRK1 -related distal Hereditary Motor Neuropathy, Bos, Rihan et al. show that the mutations in *VRK1* affect the electrical properties of these neurons: they display defects in the initiation of the Action Potential due to a shortening of the Axonal Initial Segment.

## Introduction

Mutations in *VRK1* were first described in spinal muscular atrophy with pontocerebellar hypoplasia (SMA-PCH) (Renbaum et al., 2009). Later, mutations in *VRK1* proved to be responsible for a wide spectrum of recessive neurological diseases, usually characterized by motor peripheral neuropathy, with or without sensory abnormalities, most often also upper motor neuron involvement, and occasionally structural brain changes (El-Bazzal et al., 2019). In total, 19 mutations are now reported in *VRK1* in 26 patients from 16 families, affected with diseases described under several different clinical entities: Spinal Muscular Atrophy (SMA), distal SMA (adult-onset), Amyotrophic Lateral Sclerosis (ALS), juvenile ALS, Hereditary Motor and Sensory Neuropathy (Charcot-Marie-Tooth disease), pure motor neuropathy (distal Hereditary Motor Neuropathy (dHMN)). There is no clear phenotype-genotype correlation, however, the feature, common to all patients, is the involvement of lower motor neurons (MNs). VRK1 encodes an ubiquitously expressed, mainly nuclear, serine/threonine kinase, playing a crucial role in many cellular processes like cell division and cell cycle progression (Valbuena et al., 2008; Valbuena et al., 2011).

We recently identified two new compound heterozygous missense mutations in *VRK1*, in two siblings from a Lebanese family, affected with dHMN associated with upper motor neurons (MNs) signs (El-Bazzal *et al.*, 2019).By studying patient’s cells, including induced Pluripotent Stem cell (iPS) derived Motor Neurons (hiPSC-MNs), we have shown that the mutations in *VRK1* lead to severely reduced levels of VRK1 by impairing its stability, and to a shift of nuclear VRK1 to the cytoplasm *(El-Bazzal et al., 2019*).

This depletion of VRK1 from the nucleus alters the dynamics of coilin, a phosphorylation target of VRK1 (Sanz-Garcia et al., 2011), by reducing its stability through increased proteasomal degradation (El-Bazzal *et al.*, 2019). Coilin is the main component and scaffold protein of the Cajal Bodies (CBs), membrane-free, nuclear organelles enriched in several nuclear proteins and RNA-protein complexes, and required for splicing, ribosome biogenesis and telomere maintenance (Lafarga et al., 2017). CBs are particularly prominent in post-mitotic neurons and anomalies of CBs, suc as the ones described by us in VRK1 patients’ hiPSC-MNs(El-Bazzal *et al.*, 2019), are also hallmarks of Spinal Muscular Atrophy (SMA), another MN disease (lower MN) caused by loss or mutations in the survival MN (SMN) protein (Tapia et al., 2012).

The use of human induced pluripotent stem cells (hiPSCs) (Takahashi et al., 2007; Takahashi and Yamanaka, 2006) represent a unique opportunity to study human neuronal and glial cells affected in motor neuron diseases and Inherited Peripheral Neuropathies, while maintaining the disease-associated genetic background of the patient (Saporta et al., 2011). One particular interest is the possibility to measure the electrical activity of these neurons, as demonstrated in a few studies in hiPSC-MNs from patients with Amyotrophic Lateral Sclerosis (ALS)(Sances et al., 2016; Toli et al., 2015; Wainger et al., 2014), Spinal Muscular Atrophy (SMA) (Liu et al., 2015) and Charcot-Marie-Tooth disease (CMT) (Saporta et al., 2015). In our previous work (El-Bazzal et al., 2019), we have demonstrated that hiPSC-MNs from patients with bi-allelic mutations in *VRK1* have shorter neurite length and altered branching, consistent with a length dependent axonopathy. However, putative structural changes of the axon intial segment (AIS), the site governing the spike initiation, remain unknown.

Here, we studied the electrophysiological properties of hiPSC-MNs from patients with mutations in *VRK1* as compared to controls. We confirmed that the differentiation protocol used to obtain spinal MNs from hiPSCs induces functional MNs, which are capable of displaying sustained firing patterns. We also demonstrated that co-culture with mouse myoblasts enhance the functional maturation of hiPSC-MNs. In hiPSC-MNs from patients with VRK1 mutations, the Action Potential (AP) was smaller in amplitude and larger in duration. This modification of AP appears to result from a decrease in the Axonal Initial Segment (AIS) length. These data from human MNs indicate that mutations in VRK1 contribute to the alteration of the spiking initiation that may ultimately lead to the upper and lower motor neuron disease affecting these patients.

## Results

### Differentiation of hiPSCs into spinal Motor Neurons (hiPSC-MNs)

We observed no differences in the expression of OLIG2, HB9 and ISLET1 between patient and control hiPSC-MNs, either at DIV18 or DIV30 (**Figure 1A-C**). At DIV18 half of NF-M positive cells (NF-M +) expressed OLIG2 (50.7% for control and 48.8% for patient II.2) (**Figure 1A**), while only 41 to 46 % of NF-M + cells were positive for HB9 and ISLET1, the generic markers of spinal MNs (Figure 1A-C). At DIV30, the majority (up to 80%) of neurons (NF-M + cells) were already expressing HB9 and ISLET 1, whereas OLIG2 has decreased by half, confirming the ventral spinal cord identity and the acquisition of a “mature” MN phenotype at DIV30. Furthermore, both in control and patient, at DIV30, most “mature” (HB9+/ISLET1+) cells expressed Choline Acetyltransferase (CHAT), the rate-limiting enzyme for acetylcholine, suggesting that they are likely to be functional (**Figure 1D**).

**Figure 1.**
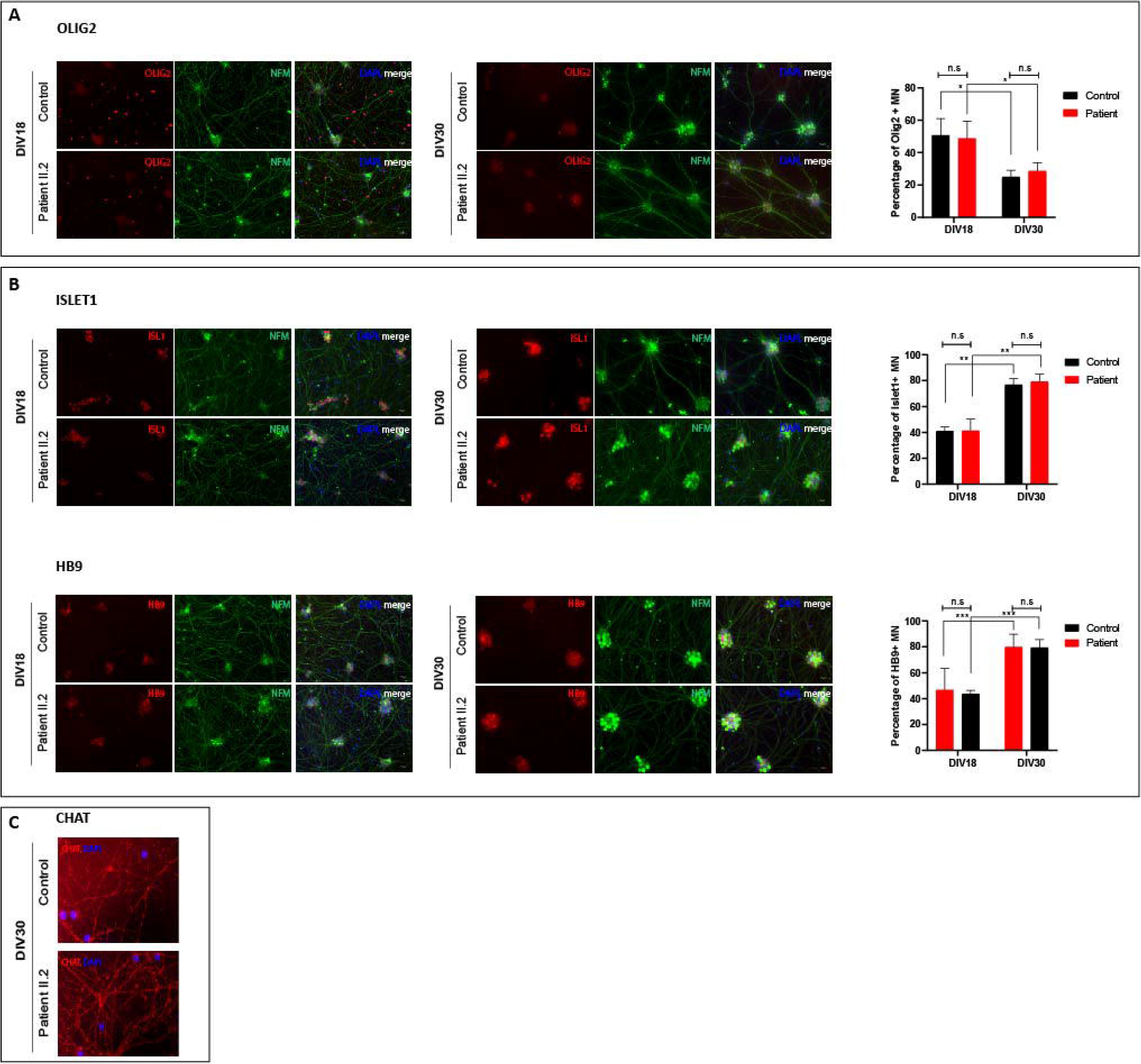
Monitoring of hiPSC differentiation to spinal Motor Neurons (hiPSC-MNs) **A.** *(Left Panel)* Immunolabeling of OLIG2, oligodendrocyte lineage transcription factor 2, a MN progenitor marker, in hiPSC-MNs from patient II.2 and control showing a decrease in OLIG2+ cells at differentiation day 30 (DIV30) as compared to differentiation day 18 (DIV18) in both individuals. NF-M: neurofilament medium chain. *(Right Panel)* Quantification of OLIG2+ neurons (NF-M+) was performed using data from three independent experiments. For each experiment, a mean of 432 and 602 cells at DIV18, and 521 and 584 cells at DIV30, were counted in the control and patient II.2 respectively. The data show a significant decrease of 41% and of 51% of OLIG2+ neurons between DIV18 and DIV30 in patient II.2’s and control’s hiPSC-MNs respectively. Statistical significance was evaluated by a two-way ANOVA test, *P<0.05, n.s: not significant. **B.** *(Left Panels)* Immunolabeling of transcription factors ISLET1 and HB9 in hiPSC-MNs from patient II.2 and a control showing an increase in HB9 +/ ISLET1 + cells at differentiation day 30 (DIV30) as compared to differentiation day 18 (DIV18) in both individuals. NF-M: Neurofilament medium chain. *(Right Panels)* Quantification of ISLET1+ neurons and HB9+ neurons (NFM+) was performed, using data from three independent experiments. For ISLET1, a mean of 881 and 876 cells at DIV18, and 1342 and 1108 cells at DIV30, were counted in each experiment, in the control and patient II.2 respectively. For HB9, we counted a mean of 745 and 780 cells at DIV18 and 1541 and 1115 cells at DIV30, in each experiment, in control and patient respectively. The data show a significant increase of 48% and of 47% of ISLET1+ neurons and of 42% and of 45% of HB9+ neurons in patient II.2’s and control’s hiPSC-MNs respectively. Statistical significance was evaluated by two-way ANOVA test, ***P<0.0005, n.s: not significant **C.** Immunolabeling of choline acetyl transferase (CHAT) in hiPSC-MNs from patient II.2 and control at DIV30 suggest that hiPSC-MNs at DIV30 are functional.

### Electrophysiological properties of hiPSC-MNs with or without coculture with myoblasts

To test whether our hiPSC-MNs were functional, we performed whole-cell recordings on hiPSC-neurons (putative hiPSC-MNs). Recorded neurons were characterized as hiPSC-MNs on the basis of their large cell bodies (>20μm), their distinct processes and their hyperpolarized resting membrane potential (<-65mV) (**Figure2A**). Before DIV30, neurons did not fire repetitively (**Figure2A, left**). These immature cells displayed transient firing with few AP in response to pulse or sinusoidal depolarizing currents.

**Figure 2.**
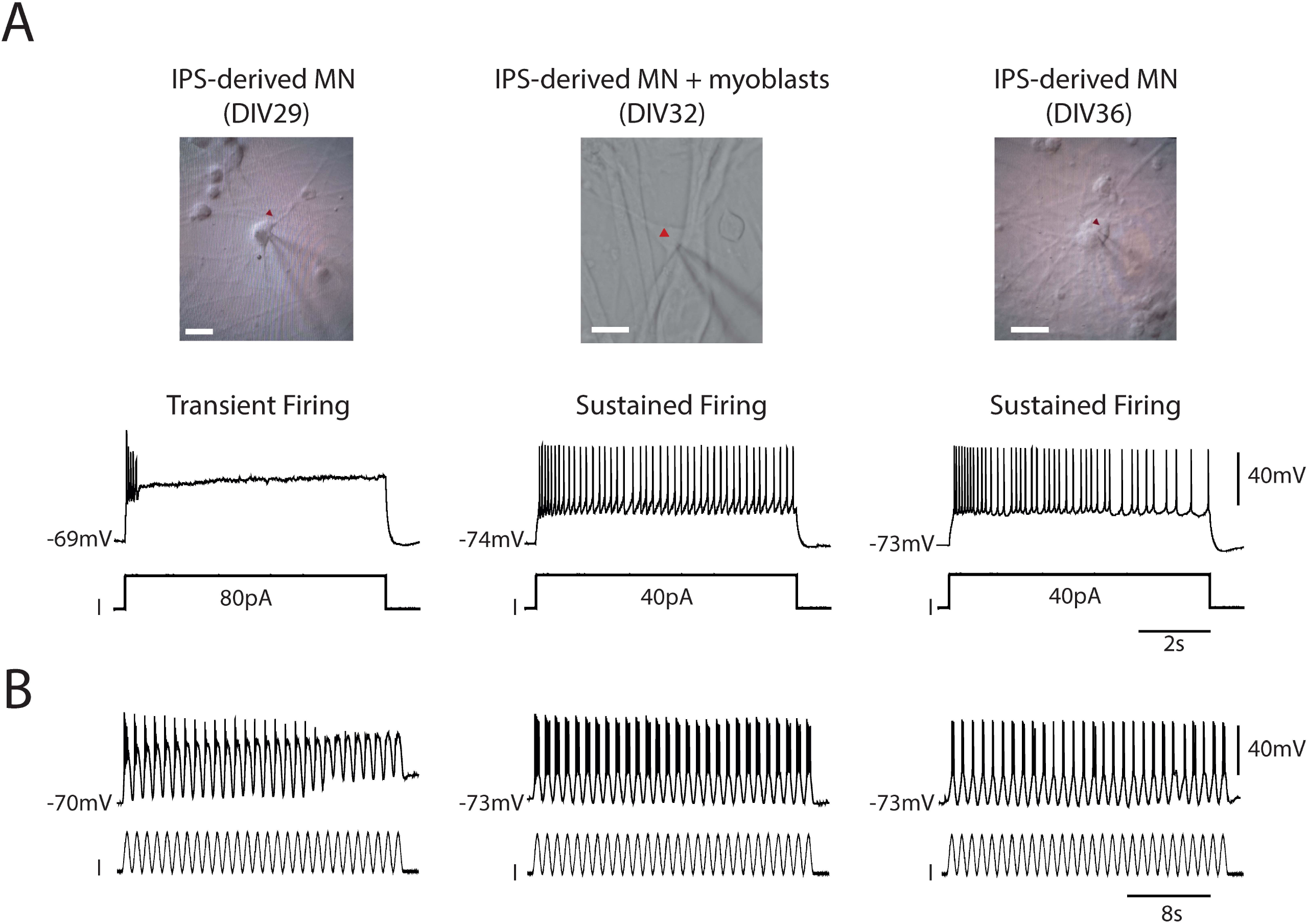
Co-culturing human hiPSC-MNs with myoblasts accelerates their functional maturation. **A.** Bright field images of hiPSC-MNs at DIV29 (left) and DIV36 (right) without co-culture of C2C12 mouse myoblasts or at DIV32 (middle) with C2C12 mouse myoblasts co-culture. The red mark indicates the recorded MNs in whole-cell configuration (current-clamp mode). Scale bar, 20uM. **B**. Representative voltage traces of three hiPS-MNs illustrated above in response to long-lasting (7.5s) depolarizing current pulses or sinusoidal current pulses (1Hz,30s).

In order to obtain further maturation of hiPSC-MNs *in vitro,* we maintained dissociated cultures for at least 2 additional days. Interestingly, we noticed that, unlike before DIV30, hiPSC-MNs from DIV32 were able to generate a robust repetitive spiking during the whole duration of the current pulse (**Figure 2A, right**). However, maintaining hiPSC-MNs in culture for more than 30 days is very challenging, because of cells detaching from their support. On the other hand, it is well known that MNs and muscle cells have mutual stabilizing and supportive influence to each other(Maffioletti et al., 2018; Mazaleyrat et al., 2020; Yoshida et al., 2015), and therefore, we sought to increase the viability and functionality of our hiPSC-MNs by coculturing them with mouse myoblasts (C2C12) for 7-9 days. This allowed obtaining large multinucleated myotubes, and hiPSC-MNs extending long neurites alongside the myotubes (**Supplementary Figure 1**). Similarly to what was observed in DIV36 hiPSC-MNs without coculture (**Figure2A-B, right**), and unlike DIV29 hiPS-MNs without coculture (**Figure 2A-B, left**), DIV32 hiPSC-MNs cocultured with myotubes (**Figure2A-B, middle**) show a mature electrophysiological phenotype and were able to sustain their firing rate without spiking inactivation in response to a prolonged depolarizing current (**Figure 2A, middle**) or in response to sinusoidal current pulses of same amplitudes (1Hz, 80pA) (**Figure 2B, middle**). This confirms the positive role of myotubes in the functional maturation of hiPSC- MNs.

### Functional subtypes diversity of hiPSC-MNs

We next tested the functional properties of mature hiPSC-MNs, either without coculture at DIV36, or at DIV32 when co-cultured for 7 days with C2C12 primary mouse myoblasts. In the latter, we specifically observed regular high-frequency contractions of myotubes (**Movie S1**). Both co-cultured and not co-cultured hiPSC-MNs display five distinct electrophysiological signatures (**Figure 3A-C**): they were able to generate three subtypes of repetitive spiking which were distinguished by a stable (**Figure 3A, left**), increased (**Figure 3A, middle**) or decreased (**Figure3A, right**) firing frequency. Recorded cells were also able to display a delay in their firing property (**Figure3B**) or a voltage-dependent oscillatory firing (**Figure3C**). For the latter subtype (**Figure3C**), increasing stimulating current intensity increased the frequency of the oscillations (gray traces). Taken together, these data indicate that the diversity of the hiPSC-MNs subtypes is a hallmark of functional maturity.

**Figure 3.**
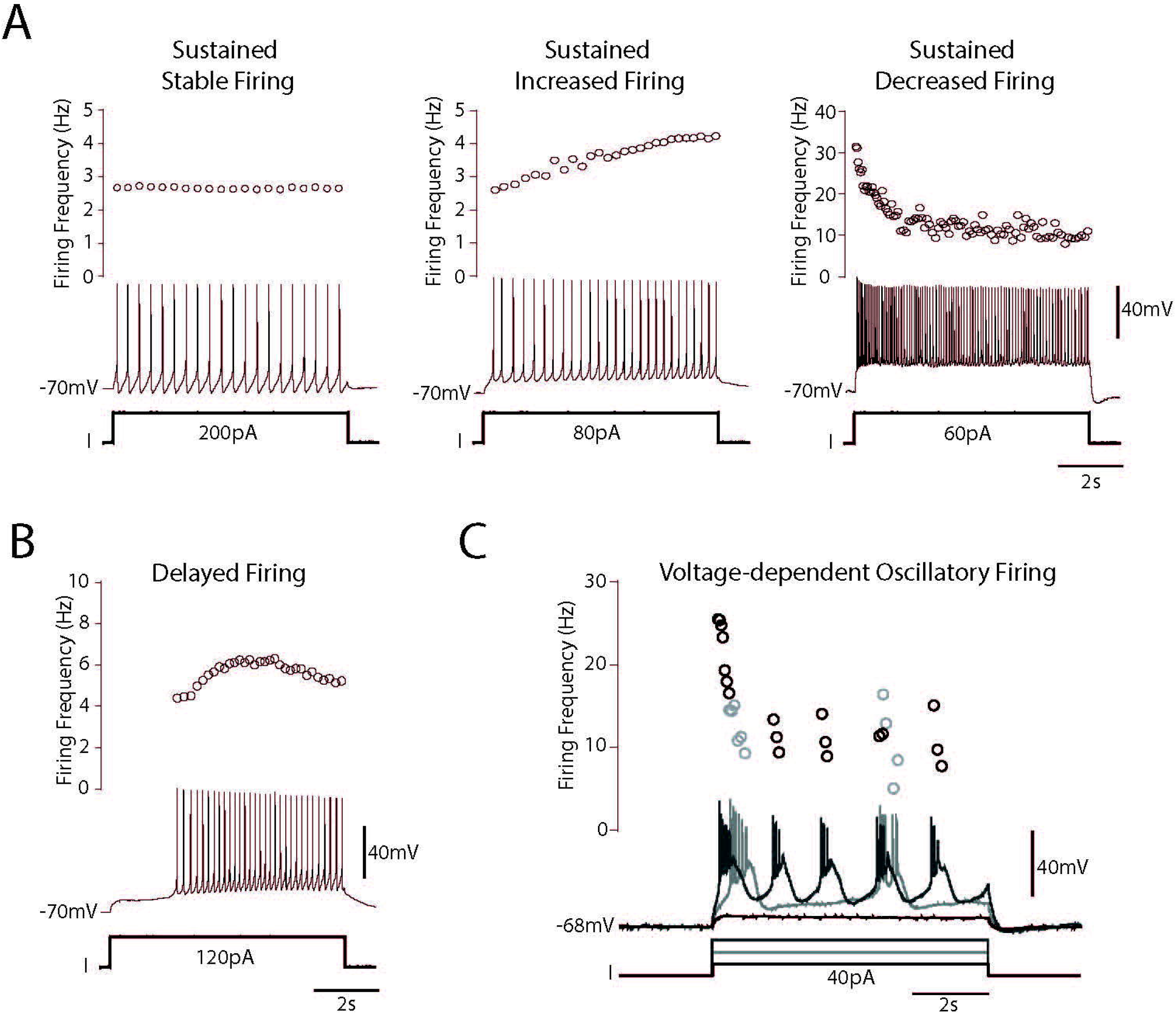
Control hiPSC-MNs co-cultured with C2C12 myoblasts display five distinct electrophysiological signatures. **A.** Representative voltage traces of three hiPSC-MNs co-cultured with C2C12 myoblasts at DIV32 displaying sustained spiking in response to long-lasting depolarizing current pulses. They are distinguished in three subtypes as a function of their firing frequency overtime: stable (left), increased (middle) or decreased (right). **B.** Representative voltage trace of one hiPSC-MN (DIV 32) co-cultured with C2C12 myoblasts displaying a delayed firing in response to a long-lasting depolarizing current pulse **C.** Representative voltage trace of one hiPS-MN co-cultured with C2C12 myoblasts at DIV32 displaying voltage-dependent oscillations in response to a long-lasting depolarizing current pulse. The frequency of the oscillations increases as a function of the current amplitude. Note the same electrophysiological patterns were recorded in IPS-derived MNs without co-culture with myoblasts.

### Electrophysiological properties of hiPSC-MNs from patients with *VRK1* mutations

In order to determine how, and if, the mutations in *VRK1* described previously by us in two patients affected with autosomal recessive distal Hereditary Motor Neuropathy (dHMN) (El-Bazzal *et al.*, 2019), affect the electrical properties of the spinal motor neurons, we recorded, in whole-cell configuration, the firing pattern of hiPSC-MNs from one patient (patient II.2, see (El-Bazzal *et al.*, 2019) from DIV29 to DIV39. We first observed that the patient’s hiPSC-MNs display the five firing patterns similar to those described in control (**Figure 4A-B**). These distinct firing subtypes were present in approximatively the same proportion than controls (**Figure 4C**). We noted only a slight decrease (3.5-5.5%) for stable and increased firing subtypes accompanied by an increase (10%) of decreased firing subtype from the patient’s hiPSC-MNs. In addition, we observed that the patient’s hiPSC-MNs were able to sustain their firing rate in response to sinusoidal current pulses of same amplitude (**Figure 4D**), similar to controls. However, we noticed a significant change in the shape of the Action Potential (AP) from the patient’s hiPSC-MNs as compared to control (**Figure 4E** and **Table 1**). In the patient’s hiPSC-MNs, we quantified a significant decrease of AP amplitude, rise slope and decay slope accompanied by an increase of AP duration and the amplitude of AHP (**Figure 4E** and **Table 1**). In contrast, the firing gain is not significantly affected in the patient (**Figure 4F**).

**Figure 4.**
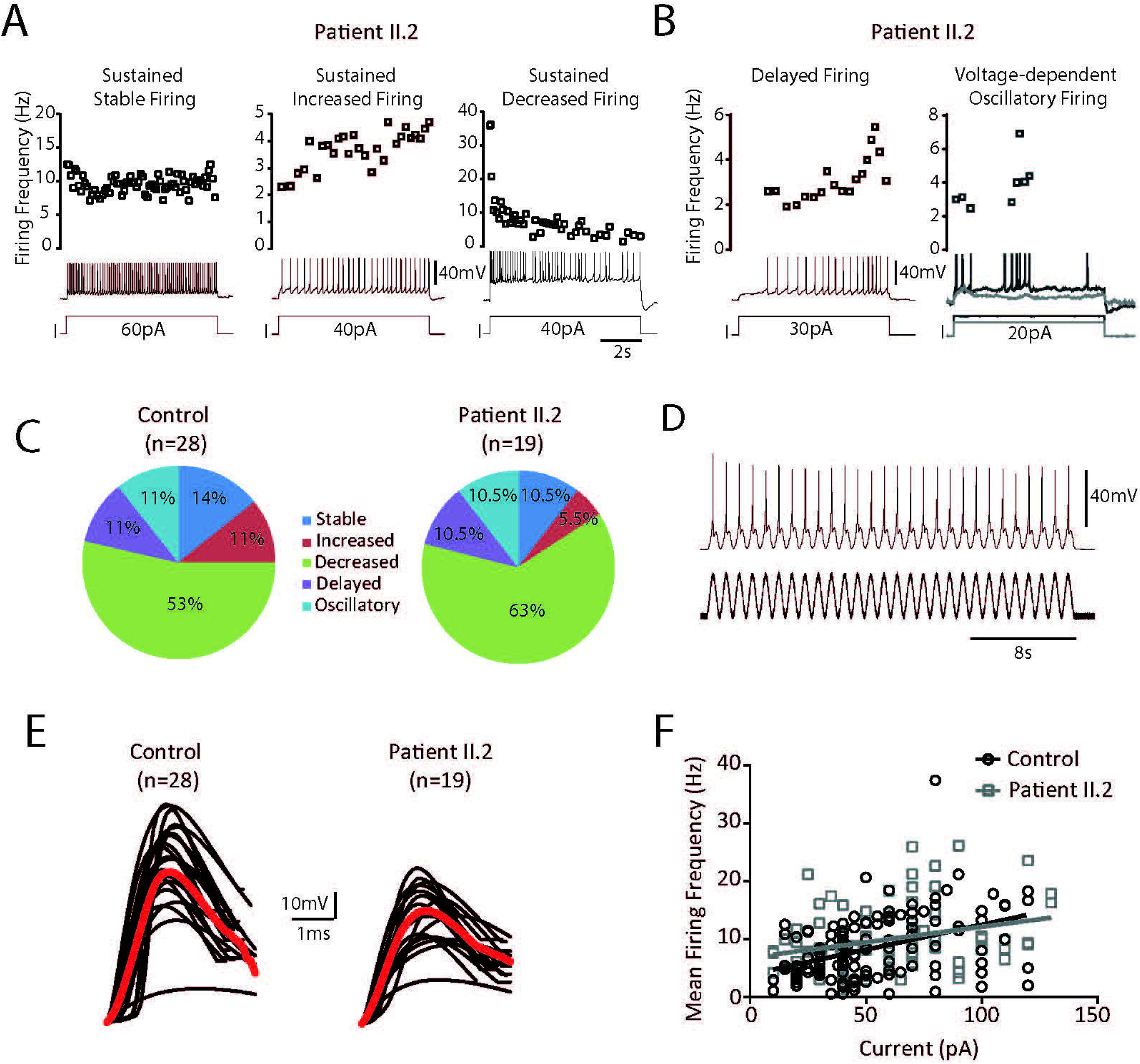
hiPSC-MNs from patient II.2 display smaller action potential. **A.** Representative voltage traces of three hiPSC-MNs from patient II.2 displaying sustained stable (left), increased (middle) or decreased (right) firing in response to long-lasting depolarizing current pulses comparable to controls at DIV32 cocultured with C2C12 myoblasts. **B.** Representative voltage traces of hiPS-MNs (DIV32, cocultured with C2C12 myoblasts) from patient II.2 displaying the delayed firing subtype (left) and the voltage-dependent oscillatory firing subtype (right) in response to a long-lasting depolarizing current pulse. **C.** Proportion of distinct firing pattern subtypes from control (left) and patient II.2 hiPSC-MNs. **D.** Representative voltage traces of three hiPSC-MNs from the patient (DIV32, cocultured with C2C12 myoblasts) displaying sustained firing in response to prolonged sinusoidal current pulse of same amplitude (1Hz, 80pA, 30s) (right). **E.** Average traces (red) of action potentials from hiPSC-MNs recorded at the rheobase from control (left) and the patient(right) at DIV32, cocultured with C2C12 myoblasts. **F.** Mean firing frequency of mature hiPSC-MNs (DIV32, cocultured with C2C12 myoblasts) as a function of injected current in control (black) and patient (gray).

**Table 1.** Action potential and firing parameters from hiPSC-MNs.

|  | Control | Patient II.2 |
| --- | --- | --- |
| n | 28 | 19 |
| AP threshold, mV | -50.33 ± 1.11 | -46.44 ± 1.21* |
| AP half width, ms | 1.1 ± 0.15 | 1.84 ± 0.39* |
| AP rise slope, mV/ms | 37.08 ± 3.76 | 23.08 ± 2.74** |
| AP decay slope, mV/ms | -12.76 ± 1.1 | -9.31 ± 1.12* |
| Peak ampl., mV | 54.01 ± 2.47 | 43.48 ± 2.07** |
| AHP ampl., mV | 9.16 ± 0.78 | 6.59 ± 0.73* |
| Rin, MΩ | 35.4 ± 4.59 | 43.06 ± 5.48 |
| Rheobase, pA | 43.04 ± 14 | 38.11 ± 4.81 |
| n | 22 | 15 |
| Gain (First ISI), Hz/pA | 0.26 ± 0.03 | 0.32 ± 0.06 |
| Gain (Steady State), Hz/pA | 0.22 ± 0.03 | 0.24 ± 0.05 |
| Gain (Whole discharge), Hz/pA | 0.22 ± 0.02 | 0.26 ± 0.05 |
| Max. Firing Rate (Hz) | 23.55 ± 2.91 | 24.73 ± 2.92 |

Data are mean±sem. \*p<0.05; \*\*p<0.01 Kruskal-Wallis test with Dunn's post-hoc test
AP: action potential; AHP: afterhyperpolarization, Rin: input resistance; ISI: inter-spike interval

### Reduced Axonal Initial (AIS) length in hiPSC-MNs from a patient with *VRK1* mutations

Previous studies performed in patient II.2’s hiPSC-MNs have shown impaired neurite length and branching in patient’s hiPSC-MNs as compared to a control(El-Bazzal *et al.*, 2019). Knowing that the AP waveform is closely linked to the Axonal Initial Segment (AIS) length (Kaphzan et al., 2011; Kuba, 2012), we sought to investigate if the modified AP amplitude observed in patient’s MNs is linked to a modification in the AIS of those MNs. To do so, we have performed immunolabeling of AnkyrinG, a scaffold protein, known as the master organizer of the AIS, in patient’s and control’s hiPSC-MNs (**Figure 5A**). While we showed no differences in the intensity of the AIS (**Figure 5C**), we observed a significant decrease in AIS length (~40%) in the patient’s MNs as compared to the control (**Figure 5B**).

**Figure 5.**
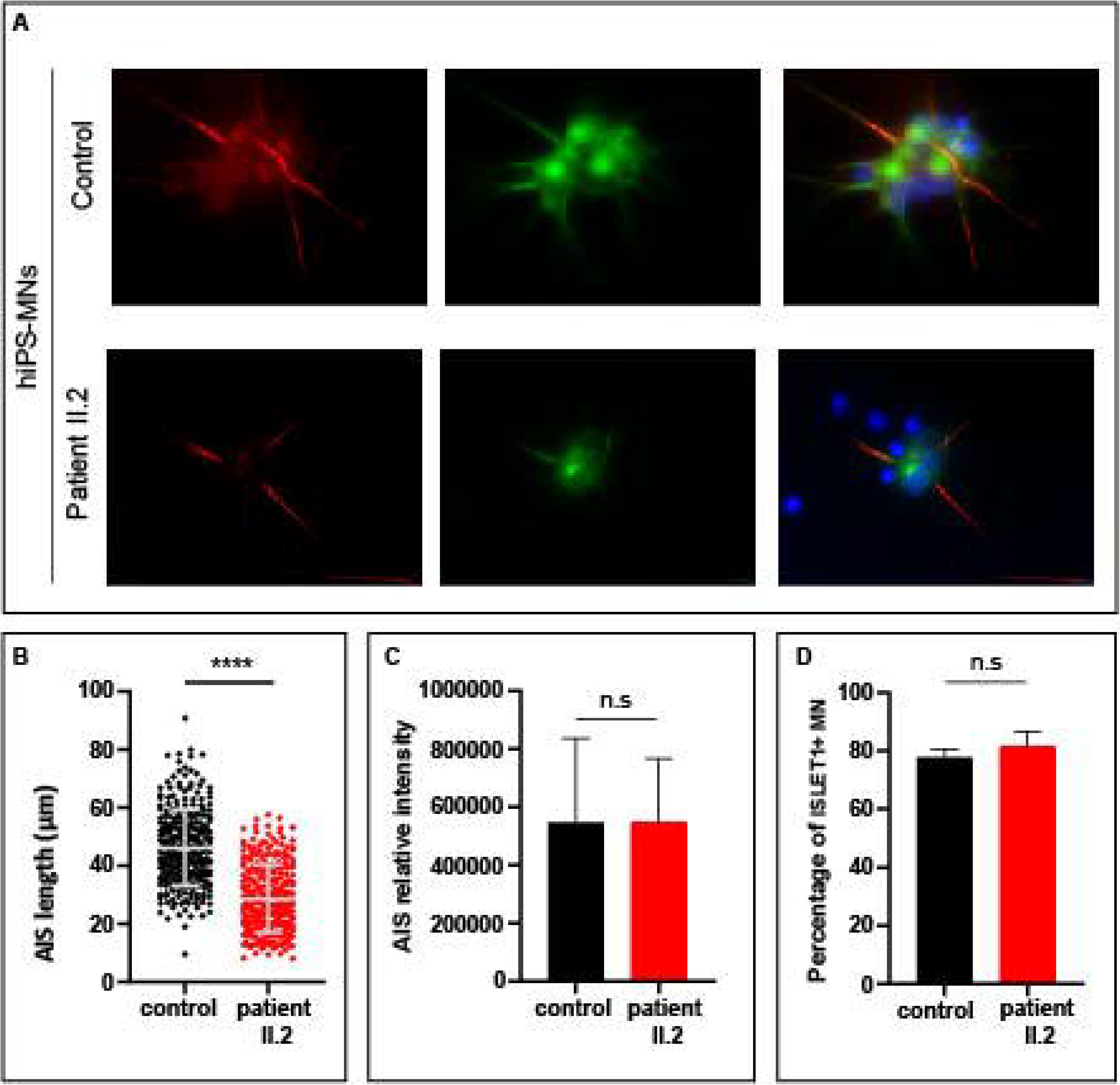
Shorter AIS in hiPSC-derived MNs from patient with *VRK1* mutations. **A**. Immunolabeling of Ankyrin G and MAP2 in hiPSC-MNs from patient II.2 as compared to control’s hiPSC-derived MNs. **B**. Measure of the AIS length in hiPSC-MNs from patient II.2 as compared to control. AIS length was defined by the length of the AnkyrinG+ segment. Measured AIS length was 46.1 μm and 28.5 μm in control and patient II.2 respectively (average of three independent experiments), showing a decrease of 38% in length in patient’s hiPSC-MNs. Measures were performed on 280 hiPSC-MNs counted from three independent experiments. Statistical significance was evaluated by Mann–Whitney test, *P <0.1, **P <0.01, ***P<0.001. **C**. Quantification of AIS intensity by measuring relative intensity of a specified area, using ImageJ software. Data were recorded, for each individual, on 75 AIS segments from three independent experiments. We did not observe any significant differences between patient II.2 and control. Statistical significance was evaluated by two-way ANOVA test: n.s: not significant. **D**. Percentage of mature hiPSC-MNs at the end of the differentiation protocol (DIV30) was evaluated by counting ISLET1 positive cells, in at least 2000 cells from three independent experiments (2007 cells for control and 2264 cells for patient II.2). These experiments are the same ones used in A-C. Statistical significance was evaluated by unpaired t-test, n.s: not significant.

## Discussion

In this study, we showed that MNs derived from human iPSCs obtained from healthy individuals or patients harbouring *VRK1* mutations develop similar electrophysiological firing subtypes. However, patient hiPSC-MNs display a shorter and larger AP in amplitude and time, respectively. This modification is likely to result from a decrease in the AIS length. These data from human MNs indicate that mutations in *VRK1* contribute to the alteration of the action potential waveform that may ultimately lead to the upper and motor neuron disease affecting these patients.

We recently described new mutations in *VRK1*, the vaccinia-related kinase 1 gene, in patients affected with a distal form of Charcot-Marie-Tooth disease (dHMN, for distal Hereditary Motor Neuropathy) associated with upper motor neuron signs (El-Bazzal *et al.*, 2019). Mutations in the *VRK1* gene are responsible for a wide spectrum of autosomal recessive neurological diseases, ranging from sensory-motor neuropathy (Charcot-Marie-Tooth disease)(Gonzaga-Jauregui et al., 2013) to juvenile ALS (Nguyen et al., 2015). All these diseases have in common the involvement of lower MNs. *VRK1* encodes the Vaccinia Related kinase 1, a 396 amino acid (aa), ubiquitously expressed, serine/threonine kinase, playing a crucial role in regulating cell cycle (Valbuena *et al.*, 2008; Valbuena *et al.*, 2011). VRK1 is mainly a nuclear protein, although the presence of a small fraction in the cytoplasm and membrane compartments has been described (Nichols and Traktman, 2004; Valbuena et al., 2007). Several substrates are known to date for VRK1(El-Bazzal *et al.*, 2019), including VRK1 (Lopez-Borges and Lazo, 2000), and coilin, the main component of Cajal Bodies (CBs)(*Sanz-Garcia et al., 2011*).

Although it is now evident that the motor tract is mainly affected in patients with mutations in *VRK1*, little is known whether these mutations directly alter the functional properties of MNs. We previously demonstrated, in human patient’s iPS-derived MNs (hiPSC-MNs), that mutations in *VRK1* cause a drop in VRK1 levels leading to Cajal bodies (CBs) disassembly and to defects in neurite outgrowth and branching (El-Bazzal *et al.*, 2019). We assumed that these changes were likely to drive a modification in cell excitability (Gulledge and Bravo, 2016). Therefore, here, we tried to answer to the question whether these structural changes affect the firing pattern of hiPSC-MNs from patients with mutations in VRK1, by performing elecrophysiological measurements in hiPSC-MNs from patients. First, we confirmed that the differentiation protocol used to obtain spinal MNs from hiPSCs (Maury et al., 2015) induces functional MNs, by testing the capacity of control hiPSC-MNs to spike in response to the injection of a current depolarizing step or repetitive oscillations. Our hiPSC-MNs showed mature phenotypes characterized by sustained firing pattern), typical of spinal motoneurons (Bos et al., 2018; Durand et al., 2015; Gao and Ziskind-Conhaim, 1998; Smith and Brownstone, 2020; Vinay et al., 2000). This sustained firing pattern occurs ~ 4-5 weeks post-plating as it has been previously shown in cultured human iPSC-derived MNs (Liu *et al.*, 2015; Maury *et al.*, 2015; Saporta *et al.*, 2015), confirming the functionality of the recorded hiPSC-MNs.

We also demonstrated that hiPSC-MNs co-cultured with mouse myoblasts enhance their functional maturation by displaying sustained firing ~ one week earlier than MNs cultures alone. This result suggests the presence of bidirectional interactions with muscle cells (Sances *et al.*, 2016) (Bucchia et al., 2018) confirmed by the presence of muscle contractions when hiPSC-MNs were firing. Because spinal motor neurons display distinct sustained firing pattern (Durand *et al.*, 2015; Hadzipasic et al., 2014; Manuel and Zytnicki, 2019), we tested whether hiPSCs-MNs also display this heterogeneity of activity pattern. Thus, we were able to clearly demonstrate the presence of distinct subtypes in sustained firing hiPSC-MNs co-cultured with muscle cells suggesting that our recorded hiPSC-MNs are fully mature. This cue of functional maturity was crucial to obtain for comparing hiPSC-MNs from healthy individuals with those from patients harbouring VRK1 mutations.

Surprisingly, although our patients’ hiPSC-MNs show structural changes, such as shorter neurite length and altered branching(El-Bazzal *et al.*, 2019), we did not observe any differences in the firing patterns of hiPSC-MNs from patients harbouring VRK1 mutations compared to healthy individuals. We, however, demonstrated significant alterations of the shape of individual action potentials, in patients, distinguished by a smaller amplitude and a larger duration. These results are in line with other studies reporting significant changes in the electrophysiological properties of hiPSC-MNs from patients affected with other diseases affecting axons of lower motor neurons (Charcot-Marie-Tooth, CMT2E and CMT2A)(Saporta *et al.*, 2015), lower motor neurons (Spinal Muscular Atrophy)(Liu *et al.*, 2015) or upper and lower motor neurons (Amyotrophic Lateral Sclerosis due to SOD1 or C9ORF72 or TARDBP 2)(Devlin et al., 2015; Wainger *et al.*, 2014).

Indeed, in those studies, all recorded hiPSC-MNs are transiently or chronically hyperexcitable, with hyperpolarized thresholds and larger AP amplitude, reflecting changes in sodium, calcium or potassium channel dynamics. In our case, hiPSC-MNs from patients displayed a depolarized AP threshold and smaller AP amplitude reflecting hypo-excitable state instead of hyperexcitability, although we did not observe any significant modification of firing gain or rheobase. Note that hypoexcitability has recently been proposed to reflect MN firing defects in mouse models of ALS(Filipchuk et al., 2021; Martinez-Silva et al., 2018), in accordance with results obtained in hiPSC-MNs of ALS patients with *C9ORF72* triplet expansions (Sareen et al., 2013). Together with our results, this underlines the need to further explore the mechanistic links between hyperexcitability and hypoexcitability.

The smaller and broader AP recorded in patients with VRK1 mutations is associated with a decrease in the length of the ankyrinG (AnkG)-positive Axonal Initial Segment (AIS). Interestingly, it has been shown that mutant mice lacking AnkG in cerebellar cells develop motor troubles and deficits in initiating AP (Zhou et al., 1998), likely due to decrease of voltage-dependent sodium channels(Jenkins and Bennett, 2001). Knowing that the AIS represents the final locus of synaptic interaction and, therefore, is an ideal target for regulating action potential initiation, AIS defects may, in turn, alter the shape of the AP. However, here, can we correlate the decrease in AIS length with the modified AP shape observed in our patient? Although only two studies does not find any positive correlation between AIS length and the spike properties (Bonnevie et al., 2020; Moubarak et al., 2019), several other studies demonstrated a clear link between these two parameters in different cell types (Gulledge and Bravo, 2016; Jorgensen et al., 2020; Kaphzan et al., 2011; Kuba et al., 2010). Indeed, Kuba *et al.*(Kuba et al., 2010) showed that auditory deprivation increased AIS length in an avian brainstem auditory neuron resulting in a bigger and shorter AP in amplitude and duration, respectively. Also, in hippocampal pyramidal neurons from a mouse model of Angelman syndrome, Kaphzan et al(Kaphzan *et al.*, 2011) demonstrate that neurons from mutant mice have longer AIS resulting in increased AP amplitude, and that AnKG and Nav1.6 are upregulated (Kaphzan *et al.*, 2011). Interestingly, in both studies(Kaphzan *et al.*, 2011; Kuba *et al.*, 2010), increased AIS length correlates with larger AP amplitude and shorter AP duration, resulting in membrane hyperexcitability, while we observe the exact opposite situation: decreased AP amplitude, longer AP duration, shorter AIS length and no hyperexcitability, as exemplified by the fact that the firing gain does not tend towards an hyper-excitable state in our patient’s hiPSC-MNs. In the G127X SOD1 Mouse Model of ALS, studies performed in mice before (Bonnevie *et al.*, 2020) and after (Jorgensen *et al.*, 2020) the onset of the disease show that plastic changes in the AIS occur around symptom onset. Knowing that changes in excitability depends more on the AIS location rather than its length (Kuba, 2012; Yamada and Kuba, 2016), the specific alteration of the AP with no significant firing defects observed here in patients’ MNs might be explained by a change specifically in the length of the AIS, but with preserved distance to soma. Unfortunately, we have not been able to measure this distance due to the fact that hiPSC-MNs differentiate from embryoid bodies, and that soma are therefore not isolated, making it difficult to identify precisely the exit point from the soma.

All together our results demonstrate that Motor Neurons from patients with VRK1-related motor neuron diseases display a modification in the shape of the AP and a depolarized AP threshold with no alteration of the proportion of firing patterns. Many work remains to be done in order to understand which pathophysiological mechanisms lead to these alterations of the AP shape and the AIS, in particular if there is a role of Nav1.6, the link with the loss of Cajal Bodies in patients’ MNs (El-Bazzal *et al.*, 2019), and finally, how the structural neurite defects previously described by us in the patients’ MNs (El-Bazzal *et al.*, 2019) are related to the AIS anomalies?

In conclusion, this study demonstrates that hiPSC-derived motor neurons can recapitulate key disease-related feature and constitute an excellent model to study the pathophysiological mechanisms leading to the diseases due to mutations in *VRK1*. Our study places this group of hereditary diseases affecting lower and upper motor neurons, in the expanding group of diseases presenting alterations of AIS components, such as epilepsy, neurodegenerative and neuroinflammatory diseases and psychiatric disorders(Leterrier, 2018). Our findings open new avenues for a better understanding, and perspectives therapeutics, not only for VRK1-related motor neuron diseases, but also for Amyotrophic Lateral Sclerosis and Spinal Muscular Atrophy, the latter sharing cellular defects with the disease affecting our patients.

## Experimental Procedures

### Patients’ cells and cell lines

The control hiPSC cell line was provided by MaSC, our Cell Reprogramming and Differentiation Facility at U1251/Marseille Medical Genetics (Marseille, France). This iPS cell line was obtained by reprogrammation of a commercial fibroblast cell line (FibroGRO™ Xeno-Free Human Foreskin Fibroblasts, #SCC058, Merck Millipore, Germany). The patient hiPSC cell line was obtained by reprogrammation of skin primary fibroblasts from the female patient II.2, affected with autosomal recessive distal Hereditary Motor Neuropathy (dHMN) due to bi-allelic in *VRK1* described in (El-Bazzal *et al.*, 2019). All hiPSCs included in the study are declared in a collection authorized by the competent authorities in France (declaration number DC-2018-3207).The mouse C2C12 myoblast cell line (#ATCC CRL-1772)(Yaffe and Saxel, 1977) was obtained from ATCC-LGC Standards (USA).

### Generation, culture and maintenance of hiPSCs

Reprogrammation of skin fibroblasts from patient and control into induced pluripotent stem cells (hiPSC) was performed by our Cell Reprogramming and Differentiation Facility (MaSC) at U 1251/Marseille Medical Genetics (Marseille, France), using Sendai virus-mediated transduction (Thermofisher Scientific, #A16517) or nucleofection (Amaxa, Lonza, # VAPD-1001 NHDF) of episomal vectors containing OCT3/4, SOX2, KLF4, C-MYC and shRNA against p53. hiPSCs colonies were picked about two weeks after transduction/nucleofection based on ES cell-like morphology. Colonies were grown and expanded in mTeSR1 medium (Stemcells technologies, #85850) on BD Matrigel™ (Corning, #354277) coated dishes. hiPSCs clones were fully characterized using classical protocols as described in (El-Bazzal *et al.*, 2019).

After thawing, hiPSCs were plated on Matrigel (Corning Life Sciences, USA, #354248) coated plastic dishes and maintained in mTeSR1 medium (Stemcell Technologies, Canada, #05851), which was exchanged daily.

### Differentiation of hiPSCs into spinal Motor Neurons (hiPSC-MNs)

For differentiation of hiPSCs into spinal MNs, we used an established 30 days differentiation protocol based on early activation of the Wnt signaling pathway, coupled to activation of the Hedgehog pathway and inhibition of Notch signaling (Maury *et al.*, 2015).

Briefly, at Day 0, hiPSCs were dissociated with accutase (Thermofisher Scientific, # A11105-01) when 70-90% confluent, and cultured, in suspension, in non-adherent, ultra-low attachment, 10 cm Petri dishes to form Embryoide Bodies (EB) in “N2B27” medium supplemented with ROCK inhibitor Y-27632 (Selleckchem, #51049) at 10μM, 20 μM of SB431542 (Stemgent, #04-0010), 0.1 μM LDN193189 (Stemgent, #04-0074), 3μM CHIR-99021 (Selleck, #51263) and 10 μM Ascorbic Acid (AA) (Sigma, #A 0278-100G).The “N2B27” medium is composed of equal volumes of ADMEM/F12 (Thermofisher Scientific, # 12634-028) and Neurobasal-A medium (Thermofisher Scientific, #10888-022), supplemented with N2 supplement (Thermofisher Scientific, # 17502-048), B-27 Supplement minus vitamin A (Thermofisher Scientific, #12587010), 200 mM Glutamax (Thermo Fisher Scientific, #35050-038).), 0.2 mM 2-mercaptoethanol (Thermofisher Scientific, #31350-010) and Penicillin/Streptomycin 1% (Thermofisher Scientific, #15070063).

At Day 2, EBs were transferred to a new dish and cultured in N2B27 supplemented with 20 μM of SB431542, 0.1 μM LDN193189, 3μM CHIR-99021, 10 μM AA and 100 nM of Retinoic Acid (RA, Sigma, #R2625-50MG). At Day 4, medium was changed for N2B27 supplemented with 20 μM of SB431542, 0.1 μM of LDN193189, 10 μM AA, 100 nM of RA and 500 nM of SAG (SIGMA, #566660). At Day 7, cells were cultured in N2B27 medium plus AA (10 μM), RA (100 nM) and SAG (500 nM). At Day 9, 10 μM of DAPT (Tocris, #2634) was added to the day 7 differentiation medium. At Day 11, EBs were dissociated in N2B27 medium supplemented with 10 μM ROCK inhibitor Y-27632, and then seeded on poly-L-ornithin (20 μg/ml, Sigma #P3655)/ Laminin (5 μg/ml, Thermofisher Scientific,#23017-015) or Matrigel (Corning, #354248) coated plates and cultured in N2B27 supplemented with 10 μM AA, 100 nM RA, 500 nM SAG, 10 μM DAPT, BDNF (Peprotech #450-02-10MG) and GDNF (Peprotech, #450-10-10MG), both at 10 ng/ml final concentration. At Day 14, SAG was removed, and the following days until Day 30, the medium (N2B27 supplemented with 10 μM AA, 100 nM RA, 10 μM DAPT, BDNF (10 ng/ml) and GDNF (10 ng/ml)) was changed every other day.

During the differentiation protocol, commitment to mature spinal MN identity was monitored by immunolabeling of markers characteristic of i) spinal MN progenitors (OLIG2), ii) canonical “mature” MN identity (transcription factors ISLET1 and HB9) and iii) more mature cholinergic MNs (CHAT, choline acetyl transferase, the rate-limiting enzyme for acetylcholine) (Davis-Dusenbery et al., 2014) at differentiation day 18 (DIV18) and 30 (DIV30) (**Figure 1A-C**). For each experiment, differentiation efficiency was assessed by HB9 and ISLET1 immunostaining as described (Maury *et al.*, 2015). Neurites were labelled using antibody to the neurofilament Medium chain (NF-M), a neuronal cytoskeletal protein.

### Culture of C2C12 myoblasts

C2C12 myoblasts were cultured in Dulbecco’s modified Eagle’s medium (DMEM) (Thermofisher Scientific, #31966021) supplemented with 10% fetal bovine serum (ThermoFisher Scientific, #10270098) and a mix of antibiotics (penicillin, streptomycin) and antimycotic (Amphotericin B) from ThermoFisher Scientific (antibiotic-antimycotic # 15240062) in a 37°C incubator stabilized at 5% CO2 until day of co-culture.

### Co-cultures of hiPSC-MNs and C2C12 myoblasts

At 7 days of culture (DIV7), C2C12 myoblasts were detached by trypsinization and 100 000 cells were plated at the top of hiPSC-MNs at day 25 of differenciation (DIV25) cultured on Matrigel-coated 6-well plates or 35 mm dishes, as described in the “Differentiation of hiPSCs into spinal Motor Neurons” section. Co-cultured cells were then maintained, during 7 days, in a medium composed of equal volumes of modified C2C12 differentiation medium (DMEM, 2% horse serum (Sigma-Aldrich, #H1270),1% antibiotic-antimycotic (ThermoFisher Scientific, #15240062) and Day14-Day30 Motor Neuron differentiation medium (see previous section). This allowed toreach 14 days of differentiation for the myotubes and 30 days for the hiPSC-MNs.

### In vitro electrophysiological recordings of hiPSC-MNs

Electrophysiological data were acquired and digitized at 4 kHz through a Digidata 1440a interface using Clampex 10 software (Molecular Devices). hiPSC-MNs were visualized with infrared differential interference contrast microscopy using a Nikon Eclipse E600FN upright microscope coupled with a water-immersion objective (Nikon Fluor 40X/0.8W). Whole-cell patch-clamp recordings were made from hiPSC-MNs without co-cultured myoblast (DIV18-DIV39) or hiPSC-MNs cocultured with myoblasts (DIV18-DIV32) selected on the basis of their large cell bodies (>20μm), their distinct processes and their hyperpolarized resting membrane potential (<-65mV). The image was enhanced with a Hitachi KP-200/201 infrared-sensitive CCD camera and displayed on a video monitor. Whole-cell patch-clamp recordings in current-clamp mode were performed with a Multiclamp 700B amplifier (Molecular Devices). Patch electrodes (2–5 MΩ) were pulled from borosilicate glass capillaries (1.5 mm OD, 1.12 mm ID; World Precision Instruments) on a Sutter P-97 puller (Sutter Instruments) and filled with intracellular solution containing the following (in mM): 140 K^+^-gluconate, 5 NaCl, 2 MgCl2, 10 HEPES, 0.5 EGTA, 2 ATP, 0.4 GTP, pH 7.3.

In the recording chamber, hiPSC-MNs were continuously bubbled with 95% O_2_ / 5% CO_2_, heated (32-34°C) and perfused with ACSF containing the following (in mM): 120 NaCl, 3 KCl, 1.25 NaH_2_PO_4_, 1.3 MgSO_4_, 1.2 CaCl_2_, 25 NaHCO_3_, 20 D-glucose, pH 7.3-7.4.

### Immunostaining and microsocopy

At Day30 of differentiation (DIV30), hiPSC-MNs, grown on coverslips in four or six well matrigel coated plates were washed with Dulbecco’s Phosphate Buffer Saline buffer (PBS) and fixed in 4% paraformaldehyde in PBS for 15 minutes. Cells were then washed for 10 min in PBS and were permeabilized using 0.1% Triton X-100 in PBS for 10 min at room temperature. After blocking for 30 min at room temperature in PBS containing 0.1%Triton X-100 and 1% Bovine Serum Albumin (BSA), hiPSC-MNs were incubated overnight at 4°c with the primary antibodies diluted in the blocking solution. Primary antibodies used were: rabbit polyclonal to Olig-2 (Merck Millipore, #AB9610) (dilution 1:500), Mouse monoclonal to Islet-1 (Biorbyt, #Orb95122)(1:200), mouse monoclonal to HB9 (DSHB, #81.5C10)(1:40), goat polyclonal to ChAT (Merck Millipore, #AB144p)(1:200), Chicken polyclonal to NF-M (Biolegend, #PCK-593P) (1:1000), rabbit polyclonal to Dysferlin (abcam, #ab124684)(1:200), and mouse monoclonal to Ankrin-G (NeuroMab, clone N106/36) (1:500).

After 3 washes in PBS, cells were incubated for 2 hours with the appropriate secondary antibody, in the blocking solution at room temperature in the dark. The following secondary antibodies were used at 1:1000 dilution: donkey Anti-Goat IgG H&L (DyLight^®^ 594) (abcam, #ab96933), donkey Anti-Mouse IgG (DyLight^®^ 550) (abcam, #ab96876), donkey Anti-Rabbit IgG (DyLight^®^ 550) (abcam, #ab96892), goat anti-Chicken IgY H&L (Alexa Fluor^®^ 488) (abcam, # ab150169)

The samples were then mounted in Vectashield mounting medium (Vector Laboratories, #H-2000) with 100ng/ml DAPI (4,6-diamidino-2-phenylindole), coverslipped, and sealed. Digitized microphotographs were recorded using an ApoTome fluorescent microscope (ZEISS, Germany), equipped with an AxioCam MRm camera. These images were merged with the ZEN software, and were treated using ImageJ software (National Institutes of Health, MD, USA).

### Measurement of AIS length and intensity

AIS length and intensity were measured on hiPSC-MNs after AnkyrinG immunolabeling, as described in the previous paragraph. AIS length and intensity were measured using ImageJ software. AIS length was defined by the length of the AnkyrinG+ segment and AIS intensity was calculated by measuring the relative intensity of the specified AnkyrinG+ area. Data for both length and intensity were recorded from three independent experiments.

## Author Contributions

R.B performed experiments, analyzed the data, interpreted the results, and wrote the manuscript

K.R. designed the study, performed experiments, analyzed the data and contributed to the writing of the manuscript

V.D. and F.B. designed the study, analyzed the data, interpreted the results and wrote the manuscript

L.E-B. contributed to the interpretation of the results and writing of the manuscript

A.M. and R.B. contributed with materials and edited the manuscript.

N.B.M, M.B., P.Q. contributed to the interpretation of the results and edited the manuscript.

## Acknowledgments

We thank MaSC, the Cell Reprogramming and Differentiation Facility at U 1251/Marseille Medical Genetics (Marseille, France) for providing the control hiPSC line.

This work was supported by grants from the French associations: ‘Association ADN’ and the French Association against Myopathies (Association Française contre les Myopathies’, Grant # TRIM-RD). This work has also been supported by a PRESTIGE programme (PRESTIGE-2017-1-0006, PCOFUND-GA-2013-609102).

We declare no conflicts of interest

## References

Bonnevie, V.S., Dimintiyanova, K.P., Hedegaard, A., Lehnhoff, J., Grondahl, L., Moldovan, M., and Meehan, C.F. (2020). Shorter axon initial segments do not cause repetitive firing impairments in the adult presymptomatic G127X SOD-1 Amyotrophic Lateral Sclerosis mouse. Sci Rep 10, 1280. 10.1038/s41598-019-57314-w.

Bos, R., Harris-Warrick, R.M., Brocard, C., Demianenko, L.E., Manuel, M., Zytnicki, D., Korogod, S.M., and Brocard, F. (2018). Kv1.2 Channels Promote Nonlinear Spiking Motoneurons for Powering Up Locomotion. Cell Rep 22, 3315–3327. 10.1016/j.celrep.2018.02.093.

Bucchia, M., Merwin, S.J., Re, D.B., and Kariya, S. (2018). Limitations and Challenges in Modeling Diseases Involving Spinal Motor Neuron Degeneration in Vitro. Frontiers in cellular neuroscience 12, 61. 10.3389/fncel.2018.00061.

Davis-Dusenbery, B.N., Williams, L.A., Klim, J.R., and Eggan, K. (2014). How to make spinal motor neurons. Development 141, 491–501. 10.1242/dev.097410.

Devlin, A.C., Burr, K., Borooah, S., Foster, J.D., Cleary, E.M., Geti, I., Vallier, L., Shaw, C.E., Chandran, S., and Miles, G.B. (2015). Human iPSC-derived motoneurons harbouring TARDBP or C9ORF72 ALS mutations are dysfunctional despite maintaining viability. Nat Commun 6, 5999. 10.1038/ncomms6999.

Durand, J., Filipchuk, A., Pambo-Pambo, A., Amendola, J., Borisovna Kulagina, I., and Gueritaud, J.P. (2015). Developing electrical properties of postnatal mouse lumbar motoneurons. Frontiers in cellular neuroscience 9, 349. 10.3389/fncel.2015.00349.

El-Bazzal, L., Rihan, K., Bernard-Marissal, N., Castro, C., Chouery-Khoury, E., Desvignes, J.P., Atkinson, A., Bertaux, K., Koussa, S., Levy, N., et al. (2019). Loss of Cajal bodies in motor neurons from patients with novel mutations in VRK1. Hum Mol Genet 28, 2378–2394. 10.1093/hmg/ddz060.

Filipchuk, A., Pambo-Pambo, A., Gaudel, F., Liabeuf, S., Brocard, C., Patrick Gueritaud, J., and Durand, J. (2021). Early hypoexcitability in a subgroup of spinal motoneurons in superoxide dismutase 1 transgenic mice, a model of amyotrophic lateral sclerosis. Neuroscience. 10.1016/j.neuroscience.2021.01.039.

Gao, B.X., and Ziskind-Conhaim, L. (1998). Development of ionic currents underlying changes in action potential waveforms in rat spinal motoneurons. Journal of neurophysiology 80, 3047–3061. 10.1152/jn.1998.80.6.3047.

Gonzaga-Jauregui, C., Lotze, T., Jamal, L., Penney, S., Campbell, I.M., Pehlivan, D., Hunter, J.V., Woodbury, S.L., Raymond, G., Adesina, A.M., et al. (2013). Mutations in VRK1 associated with complex motor and sensory axonal neuropathy plus microcephaly. JAMA Neurol 70, 1491–1498. 10.1001/jamaneurol.2013.45981754326 [pii].

Gulledge, A.T., and Bravo, J.J. (2016). Neuron Morphology Influences Axon Initial Segment Plasticity. eNeuro 3. 10.1523/ENEURO.0085-15.2016.

Hadzipasic, M., Tahvildari, B., Nagy, M., Bian, M., Horwich, A.L., and McCormick, D.A. (2014). Selective degeneration of a physiological subtype of spinal motor neuron in mice with SOD1-linked ALS. Proc Natl Acad Sci U S A 111, 16883–16888. 10.1073/pnas.1419497111.

Jenkins, S.M., and Bennett, V. (2001). Ankyrin-G coordinates assembly of the spectrin-based membrane skeleton, voltage-gated sodium channels, and L1 CAMs at Purkinje neuron initial segments. J Cell Biol 155, 739–746. 10.1083/jcb.200109026.

Jorgensen, H.S., Jensen, D.B., Dimintiyanova, K.P., Bonnevie, V.S., Hedegaard, A., Lehnhoff, J., Moldovan, M., Grondahl, L., and Meehan, C.F. (2020). Increased Axon Initial Segment Length Results in Increased Na(+) Currents in Spinal Motoneurones at Symptom Onset in the G127X SOD1 Mouse Model of Amyotrophic Lateral Sclerosis. Neuroscience. 10.1016/j.neuroscience.2020.11.016.

Kaphzan, H., Buffington, S.A., Jung, J.I., Rasband, M.N., and Klann, E. (2011). Alterations in intrinsic membrane properties and the axon initial segment in a mouse model of Angelman syndrome. J Neurosci 31, 17637–17648. 10.1523/JNEUROSCI.4162-11.2011.

Kuba, H. (2012). Structural tuning and plasticity of the axon initial segment in auditory neurons. J Physiol 590, 5571–5579. 10.1113/jphysiol.2012.237305.

Kuba, H., Oichi, Y., and Ohmori, H. (2010). Presynaptic activity regulates Na(+) channel distribution at the axon initial segment. Nature 465, 1075–1078. 10.1038/nature09087.

Lafarga, M., Tapia, O., Romero, A.M., and Berciano, M.T. (2017). Cajal bodies in neurons. RNA Biol 14, 712–725. 10.1080/15476286.2016.1231360.

Leterrier, C. (2018). The Axon Initial Segment: An Updated Viewpoint. J Neurosci 38, 2135–2145. 10.1523/JNEUROSCI.1922-17.2018.

Liu, H., Lu, J., Chen, H., Du, Z., Li, X.J., and Zhang, S.C. (2015). Spinal muscular atrophy patient-derived motor neurons exhibit hyperexcitability. Sci Rep 5, 12189. 10.1038/srep12189.

Lopez-Borges, S., and Lazo, P.A. (2000). The human vaccinia-related kinase 1 (VRK1) phosphorylates threonine-18 within the mdm-2 binding site of the p53 tumour suppressor protein. Oncogene 19, 3656–3664. 10.1038/sj.onc.1203709.

Maffioletti, S.M., Sarcar, S., Henderson, A.B.H., Mannhardt, I., Pinton, L., Moyle, L.A., Steele-Stallard, H., Cappellari, O., Wells, K.E., Ferrari, G., et al. (2018). Three-Dimensional Human iPSC-Derived Artificial Skeletal Muscles Model Muscular Dystrophies and Enable Multilineage Tissue Engineering. Cell Rep 23, 899–908. 10.1016/j.celrep.2018.03.091.

Manuel, M., and Zytnicki, D. (2019). Molecular and electrophysiological properties of mouse motoneuron and motor unit subtypes. Current opinion in physiology 8, 23–29. 10.1016/j.cophys.2018.11.008.

Martinez-Silva, M.L., Imhoff-Manuel, R.D., Sharma, A., Heckman, C.J., Shneider, N.A., Roselli, F., Zytnicki, D., and Manuel, M. (2018). Hypoexcitability precedes denervation in the large fast-contracting motor units in two unrelated mouse models of ALS. eLife 7. 10.7554/eLife.30955.

Maury, Y., Come, J., Piskorowski, R.A., Salah-Mohellibi, N., Chevaleyre, V., Peschanski, M., Martinat, C., and Nedelec, S. (2015). Combinatorial analysis of developmental cues efficiently converts human pluripotent stem cells into multiple neuronal subtypes. Nat Biotechnol 33, 89–96. 10.1038/nbt.3049nbt.3049 [pii].

Mazaleyrat, K., Badja, C., Broucqsault, N., Chevalier, R., Laberthonniere, C., Dion, C., Baldasseroni, L., El-Yazidi, C., Thomas, M., Bachelier, R., et al. (2020). Multilineage Differentiation for Formation of Innervated Skeletal Muscle Fibers from Healthy and Diseased Human Pluripotent Stem Cells. Cells 9. 10.3390/cells9061531.

Moubarak, E., Engel, D., Dufour, M.A., Tapia, M., Tell, F., and Goaillard, J.M. (2019). Robustness to Axon Initial Segment Variation Is Explained by Somatodendritic Excitability in Rat Substantia Nigra Dopaminergic Neurons. J Neurosci 39, 5044–5063. 10.1523/JNEUROSCI.2781-18.2019.

Nguyen, T.P., Biliciler, S., Wiszniewski, W., and Sheikh, K. (2015). Expanding Phenotype of VRK1 Mutations in Motor Neuron Disease. J Clin Neuromuscul Dis 17, 69–71. 10.1097/CND.000000000000009600131402-201512000-00005 [pii].

Nichols, R.J., and Traktman, P. (2004). Characterization of three paralogous members of the Mammalian vaccinia related kinase family. J Biol Chem 279, 7934–7946. 10.1074/jbc.M310813200.

Renbaum, P., Kellerman, E., Jaron, R., Geiger, D., Segel, R., Lee, M., King, M.C., and Levy-Lahad, E. (2009). Spinal muscular atrophy with pontocerebellar hypoplasia is caused by a mutation in the VRK1 gene. Am J Hum Genet 85, 281–289. 10.1016/j.ajhg.2009.07.006S0002-9297(09)00297-3 [pii].

Sances, S., Bruijn, L.I., Chandran, S., Eggan, K., Ho, R., Klim, J.R., Livesey, M.R., Lowry, E., Macklis, J.D., Rushton, D., et al. (2016). Modeling ALS with motor neurons derived from human induced pluripotent stem cells. Nature neuroscience 19, 542–553. 10.1038/nn.4273.

Sanz-Garcia, M., Vazquez-Cedeira, M., Kellerman, E., Renbaum, P., Levy-Lahad, E., and Lazo, P.A. (2011). Substrate profiling of human vaccinia-related kinases identifies coilin, a Cajal body nuclear protein, as a phosphorylation target with neurological implications. J Proteomics 75, 548–560. 10.1016/j.jprot.2011.08.019.

Saporta, M.A., Dang, V., Volfson, D., Zou, B., Xie, X.S., Adebola, A., Liem, R.K., Shy, M., and Dimos, J.T. (2015). Axonal Charcot-Marie-Tooth disease patient-derived motor neurons demonstrate disease-specific phenotypes including abnormal electrophysiological properties. Exp Neurol 263, 190–199. 10.1016/j.expneurol.2014.10.005S0014-4886(14)00341-0 [pii].

Saporta, M.A., Grskovic, M., and Dimos, J.T. (2011). Induced pluripotent stem cells in the study of neurological diseases. Stem Cell Res Ther 2, 37. 10.1186/scrt78scrt78 [pii].

Sareen, D., O’Rourke, J.G., Meera, P., Muhammad, A.K., Grant, S., Simpkinson, M., Bell, S., Carmona, S., Ornelas, L., Sahabian, A., et al. (2013). Targeting RNA foci in iPSC-derived motor neurons from ALS patients with a C9ORF72 repeat expansion. Science translational medicine 5, 208ra149. 10.1126/scitranslmed.3007529.

Smith, C.C., and Brownstone, R.M. (2020). Spinal motoneuron firing properties mature from rostral to caudal during postnatal development of the mouse. J Physiol 598, 5467–5485. 10.1113/JP280274.

Takahashi, K., Tanabe, K., Ohnuki, M., Narita, M., Ichisaka, T., Tomoda, K., and Yamanaka, S. (2007). Induction of pluripotent stem cells from adult human fibroblasts by defined factors. Cell 131, 861–872. S0092-8674(07)01471-7 [pii] 10.1016/j.cell.2007.11.019.

Takahashi, K., and Yamanaka, S. (2006). Induction of pluripotent stem cells from mouse embryonic and adult fibroblast cultures by defined factors. Cell 126, 663–676. S0092-8674(06)00976-7 [pii] 10.1016/j.cell.2006.07.024.

Tapia, O., Bengoechea, R., Palanca, A., Arteaga, R., Val-Bernal, J.F., Tizzano, E.F., Berciano, M.T., and Lafarga, M. (2012). Reorganization of Cajal bodies and nucleolar targeting of coilin in motor neurons of type I spinal muscular atrophy. Histochemistry and cell biology 137, 657–667. 10.1007/s00418-012-0921-8.

Toli, D., Buttigieg, D., Blanchard, S., Lemonnier, T., Lamotte d’Incamps, B., Bellouze, S., Baillat, G., Bohl, D., and Haase, G. (2015). Modeling amyotrophic lateral sclerosis in pure human iPSc-derived motor neurons isolated by a novel FACS double selection technique. Neurobiol Dis 82, 269–280. 10.1016/j.nbd.2015.06.011.

Valbuena, A., Lopez-Sanchez, I., and Lazo, P.A. (2008). Human VRK1 is an early response gene and its loss causes a block in cell cycle progression. PloS one 3, e1642. 10.1371/journal.pone.0001642.

Valbuena, A., Lopez-Sanchez, I., Vega, F.M., Sevilla, A., Sanz-Garcia, M., Blanco, S., and Lazo, P.A. (2007). Identification of a dominant epitope in human vaccinia-related kinase 1 (VRK1) and detection of different intracellular subpopulations. Archives of biochemistry and biophysics 465, 219–226. 10.1016/j.abb.2007.06.005.

Valbuena, A., Sanz-Garcia, M., Lopez-Sanchez, I., Vega, F.M., and Lazo, P.A. (2011). Roles of VRK1 as a new player in the control of biological processes required for cell division. Cell Signal 23, 1267–1272. 10.1016/j.cellsig.2011.04.002.

Vinay, L., Brocard, F., and Clarac, F. (2000). Differential maturation of motoneurons innervating ankle flexor and extensor muscles in the neonatal rat. The European journal of neuroscience 12, 4562–4566. 10.1046/j.0953-816x.2000.01321.x.

Wainger, B.J., Kiskinis, E., Mellin, C., Wiskow, O., Han, S.S., Sandoe, J., Perez, N.P., Williams, L.A., Lee, S., Boulting, G., et al. (2014). Intrinsic membrane hyperexcitability of amyotrophic lateral sclerosis patient-derived motor neurons. Cell Rep 7, 1–11. 10.1016/j.celrep.2014.03.019.

Yaffe, D., and Saxel, O. (1977). Serial passaging and differentiation of myogenic cells isolated from dystrophic mouse muscle. Nature 270, 725–727. 10.1038/270725a0.

Yamada, R., and Kuba, H. (2016). Structural and Functional Plasticity at the Axon Initial Segment. Frontiers in cellular neuroscience 10, 250. 10.3389/fncel.2016.00250.

Yoshida, M., Kitaoka, S., Egawa, N., Yamane, M., Ikeda, R., Tsukita, K., Amano, N., Watanabe, A., Morimoto, M., Takahashi, J., et al. (2015). Modeling the early phenotype at the neuromuscular junction of spinal muscular atrophy using patient-derived iPSCs. Stem Cell Reports 4, 561–568. 10.1016/j.stemcr.2015.02.010.

Zhou, D., Lambert, S., Malen, P.L., Carpenter, S., Boland, L.M., and Bennett, V. (1998). AnkyrinG is required for clustering of voltage-gated Na channels at axon initial segments and for normal action potential firing. J Cell Biol 143, 1295–1304. 10.1083/jcb.143.5.1295.

