## Supplementary figures and images for "Altered action potential waveform and shorter axonal initial segment in hiPSC-derived motor neurons with mutations in *VRK1*"

### Supplemental Figure1

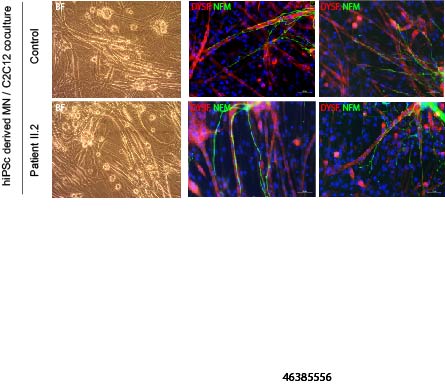
